# Mice carrying a humanized *Foxp2* knock-in allele show region-specific shifts of striatal Foxp2 expression levels

**DOI:** 10.1101/514893

**Authors:** C Schreiweis, T Irinopoulou, B Vieth, L Laddada, F Oury, E Burguière, W Enard, M Groszer

**Affiliations:** Inserm UMR-S 839, 75005 Paris, France; Sorbonne Université, 75005 Paris, France; Institut du Fer à Moulin, 75005 Paris, France; Anthropology and Human Genomics, Department of Biology II, Ludwig Maximilians University Munich, D-82152 Martinsried, Germany; Genetics, Reproduction and Development Laboratory (GReD) Genetics, Reproduction and Development Laboratory, INSERM U1103, CNRS UMR6293, Clermont-Auvergne University, Clermont-Ferrand, France; Institut Necker Enfants Malades, INSERM UMR_S1151 CNRS UMR8253, Université Paris Descartes F-75014 Paris, France; Sorbonne Université, INSERM U 1127, CNRS UMR 7225, Institut du Cerveau et de la Moelle épinière, F-75013 Paris, France

**Keywords:** Foxp2, striatum, expression, striosome, evolution

## Abstract

Genetic and clinical studies of speech and language disorders are providing starting points to unravel underlying neurobiological mechanisms. The gene encoding the transcription factor *FOXP2* has been the first example of a gene involved in the development and evolution of this human-specific trait. A number of autosomal-dominant *FOXP2* mutations are associated with developmental speech and language deficits indicating that gene dosage plays an important role in the disorder. Comparative genomics studies suggest that two human-specific amino acid substitutions in FOXP2 might have been positively selected during human evolution. A knock-in mouse model carrying these two amino acid changes in the endogenous mouse *Foxp2* gene *(Foxp2*^*hum/hum*^) shows profound changes in striatum-dependent behaviour and neurophysiology, supporting a functional role for these changes. However, how this affects Foxp2 expression patterns in different striatal regions and compartments has not been assessed. Here, we characterized Foxp2 protein expression patterns in adult striatal tissue in *Foxp2*^*hum/hum*^ mice. Consistent with prior reports in wildtype mice, we find that striatal neurons in *Foxp2*^*hum/hum*^ mice and wildtype littermates express Foxp2 in a range from low to high levels. However, we observe a shift towards more cells with higher Foxp2 expression levels in *Foxp2*^*hum/hum*^ mice, significantly depending on the striatal region and the compartment. As potential behavioural readout of these shifts in Foxp2 levels across striatal neurons, we employed a morphine sensitization assay. While we did not detect differences in morphine-induced hyperlocomotion during acute treatment, there was an attenuated hyperlocomotion plateau during sensitization in *Foxp2*^*hum/hum*^ mice. Taken together, these results suggest that the humanized *Foxp2* allele in a mouse background is associated with a shift in striatal Foxp2 protein expression pattern.

## INTRODUCTION

Genetic studies of rare and common disorders affecting speech and/or language development have provided starting points to unravel the biological basis of this human specific trait (Deriziotis & Fisher, 2017). The gene encoding the transcription factor forkhead box P2 (*FOXP2*) (for *FOXP2* nomenclature see methods part) is the first example for this approach. Humans with only one functional copy of *FOXP2* experience difficulties in learning and performing complex orofacial movements and have wide-ranging receptive and expressive deficits in oral and written language (Watkins, Dronkers, & Vargha-Khadem, 2002a).

This FOXP2 dosage deficit is thought to precipitate at cortico-basal ganglia and cortico-cerebellar neural circuits (Liégeois et al., 2003; Vargha-Khadem et al., 1998; Vargha-Khadem, Gadian, Copp, & Mishkin, 2005; Watkins, Vargha-Khadem, et al., 2002b). These circuits are essential for a number of cognitive and motor functions including learning, reward signaling and automatisation of thoughts and movements (Graybiel, 2008)

Various vertebrate species show a conserved coding sequence and similar Foxp2 expression patterns in homologous brain regions, prominently in cortex, striatum, thalamus and cerebellum (Ferland, Cherry, Preware, Morrisey, & Walsh, 2003; Rodenas-Cuadrado et al., 2018; Takahashi, Liu, Hirokawa, & Takahashi, 2003; Takahashi et al., 2008). This has led to study Foxp2 functions in animal models to delineate mechanisms potentially recruited and adapted in human evolution. For instance in songbirds, *FoxP2* mRNA is acutely downregulated within a dedicated cortico-striatal nucleus (Area X) when adult males sing (Teramitsu & White, 2006). Experimental reduction of *FoxP2* levels in Area X causes impaired song acquisition potentially due to deficits in dopaminergic signalling modulation (Haesler et al., 2007; Murugan, Harward, Scharff, & Mooney, 2013). In mice about 95% of striatal neurons comprise medium spiny neurons (MSNs), approximately half expressing the dopamine D1 receptor (D1R) the other half dopamine D2 receptor (D2R) (Kreitzer, 2009). Foxp2 expression has been found to be generally high in D1R MSNs and low in D2R MSNs (Heiman et al., 2008; Lobo, Karsten, Gray, Geschwind, & Yang, 2006; Vernes et al., 2011), indicating a range of expression levels. Foxp2 haploinsufficiency in mice has been associated with deficits in motor learning and related striatal synaptic plasticity (French et al., 2012; Groszer et al., 2008; Shu et al., 2005).

When comparing human proteins to orthologs in mouse, birds, frog and fish, FOXP2 is amongst the 5% most conserved proteins (Enard, 2011). During six million years of human evolution, two amino acid substitutions have occurred in the transcription factor encoded by *FOXP2* (Fig.1A), more than expected given its conservation in primates and mammals (Enard, 2011; Enard et al., 2002; Zhang, Webb, & Podlaha, 2002). These two substitutions happened on the human lineage after the split from the most recent common ancestor with the chimpanzee (Enard et al., 2002). Both amino acid differences localize outside the DNA binding domain in exon 7 of the *FOXP2* gene and may be implicated in protein-protein interactions (Enard et al., 2002).

**Figure 1.**
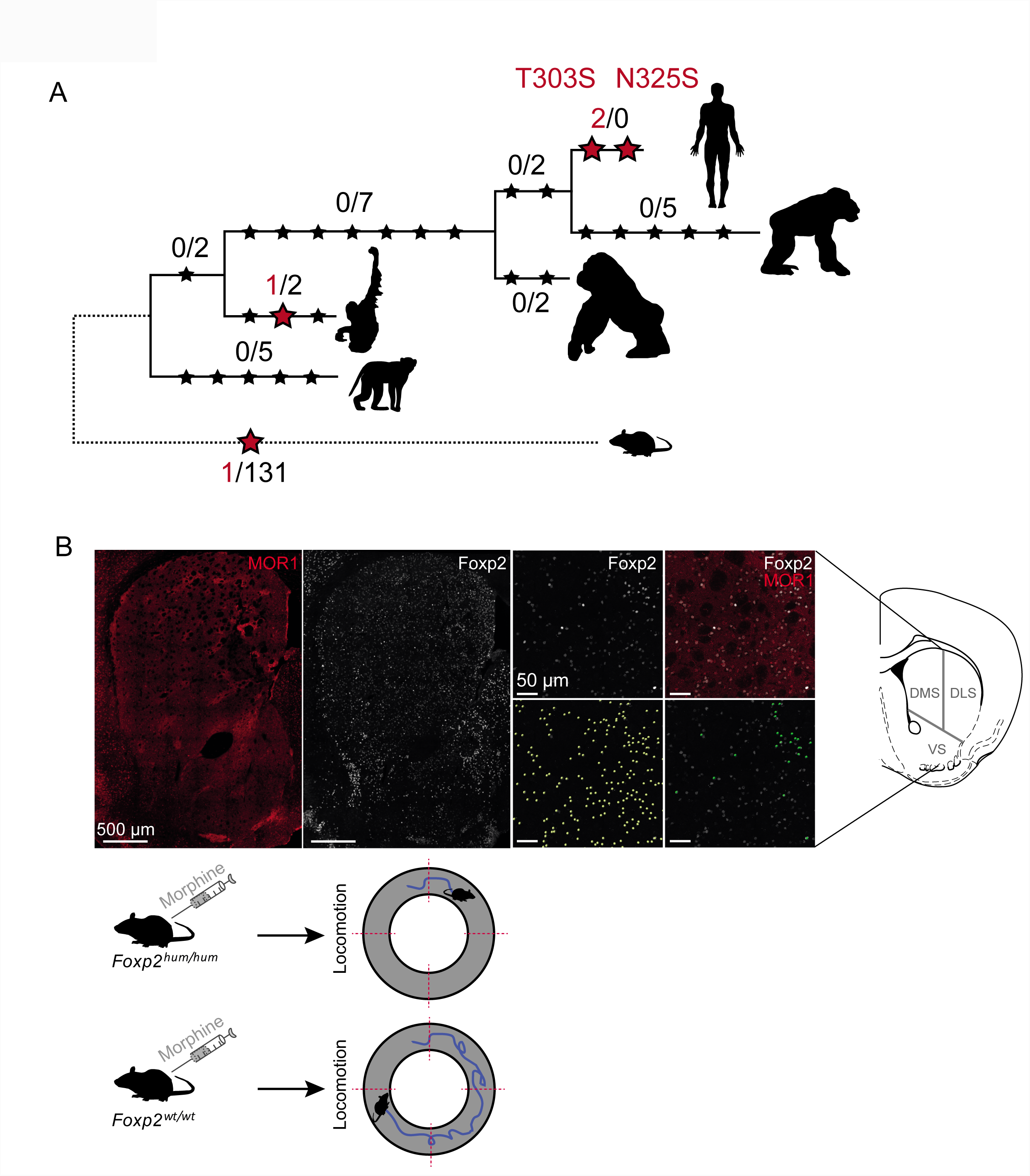
Overview of study design. (A) FOXP2 is a highly conserved transcription factor that changed at two positions during human evolution. This is more than expected given the number of silent substitutions (black stars) and number of amino acid substitutions (red stars) on other branches. (B) Knock-in mice homozygous for a *Foxp2* allele that carries these two amino acid changes (*Foxp2*^*hum/hum*^) in comparison to their wildtype littermates serve as a model to study the effects of these two amino acid changes. In this study we compared these mice for the density of Foxp2-positive neurons in three striatal regions (ventral, dorsomedial and dorsolateral) located in the matrix (MOR1-) and striosomal compartment. Left histology image panels show an exemplary coronal section of adult wildtype mouse striatum, stained either for MORI (red, left) or Foxp2 (white, right). Right histology panels show a magnification of this tissue example with histochemically labelled Foxp2 (white) or double-labelled for MORI (red) / Foxp2 (white), as well as the automatically quantified total amount of Foxp2 signal (yellow dots) as well as the highly Foxp2+ signal (green dots) in the bottom panels. We furthermore compared the locomotor activity of *Foxp2*^*hum/hum*^ and *Foxp2*^*wt/wt*^ mice in a morphine sensitization assay.

Initial genomic studies in humans detected a significantly reduced frequency of intronic DNA polymorphisms upstream of exon 7. It suggested that this genome region swept in the relatively short evolutionary timeframe of less than 200kyrs throughout the human population (selective sweep) due to strong positive selection of a beneficial mutation. (Enard et al., 2002; Zhang, Webb, & Podlaha, 2002). However, subsequently it was found that Neanderthals (Green et al., 2010; Krause et al., 2007) and Denisovans (Reich et al., 2010) also have these two amino acid substitutions and that the haplotype structure across exon 7 in currently living humans is also not compatible with a selective sweep that is caused by the two amino acid substitutions (Ptak et al., 2009; Maricic et al., 2013). Recently, the sweep signal was re-evaluated in hundreds of globally distributed genomes and while the original signal could be reproduced, it is more likely to be caused by the particular sample composition in the original data than by a selective sweep (Atkinson et al., 2018). Indeed, it has become clear that the power to detect any selective sweep that happened before humans migrated out of Africa is very low (Huber, DeGiorgio, Hellmann, & Nielsen, 2016; Nielsen, Hellmann, Hubisz, Bustamante, & Clark, 2007). Thus while the sweep signal and the amino acid substitutions are independent, the role of FOXP2 in speech and language development and the two amino acid changes argue for its role in speech and language evolution (Enard, 2014, 2016). Further support came from functional studies in mice, carrying ‘humanized’ *Foxp2* alleles (*Foxp2*^*hum/hum*^ *mice)*. *Foxp2*^*hum/hum*^ mice show striatal abnormalities in neuromorphological, neurochemical and neurophysiological parameters and behavioural effects related to cortico-basal ganglia circuits (Enard et al., 2009; Reimers-Kipping, Hevers, Pääbo, & Enard, 2011; Schreiweis et al., 2014). Specifically, we observed an accelerated transition from declarative to procedural learning in *Foxp2*^*hum/hum*^ mice as compared to wildtype littermates. This is correlated with a shifted balance of dopamine tissue levels and dopamine-dependent synaptic plasticity between the dorsomedial and dorsolateral striatal regions in *Foxp2*^*hum/hum*^ mice (Schreiweis et al., 2014).

The histoarchitecture of the striatum has been traditionally classified into regions and compartments based on afferent and efferent connectivity and the expression of specific genes (Kreitzer, 2009). During development, Foxp2 has been prominently detected in the so-called striosome compartment (Chen et al., 2016; Takahashi et al., 2003, 2008), which occupies around 10-15% of the striatal volume. It intersperses all striatal subregions as a three-dimensional, ‘labyrinthine’-like network of clustered cell-islands and is traditionally characterized mainly by histochemical markers such as μ-opioid receptor 1 (MOR1) or substance P (Brimblecombe & Cragg, 2017; Crittenden & Graybiel, 2011; Gerfen & Bolam, 2010; Graybiel & Ragsdale, 1978). Striosomal neurons are born before matrix neurons migrate more freely in between the more stationary striosomal neurons in a second developmental wave (Hagimoto, Takami, Murakami, & Tanabe, 2017; Song & Harlan, 1994; van der Kooy & Fishell, 1987), creating an adult striatal mosaic. One exception to this developmental pattern is the observation that striosomes of the ventral striatum are formed during a time point that presumably overlaps with the formation of certain matrix neurons dorsally (Song & Harlan, 1994). Cortico-striatal input of striosomes versus matrix is topographically organized and predominantly originates from limbic versus motor areas, respectively (Alexander, DeLong, & Strick, 1986; Crittenden & Graybiel, 2011; Gerfen, 1989; Gerfen & Bolam, 2010). The probably unique hodological feature of striosomes is their direct projection onto dopaminergic neurons in the midbrain (Crittenden & Graybiel, 2011; Gerfen & Bolam, 2010; Watabe-Uchida, Zhu, Ogawa, Vamanrao, & Uchida, 2012). While development and structure of the striosome-matrix compartmentalization are increasingly well understood, the physiological characterization is just beginning. Recent studies show a differentiated involvement of striosome-matrix compartments in substance P-mediated dopamine release, as well as a different modulation across reward-conditioned learning (Bloem, Huda, Sur, & Graybiel, 2017; Brimblecombe & Cragg, 2015; Yoshizawa, Ito, & Doya, 2018).

While loss of function investigations in humans, mice and songbirds have pointed to a prominent role of Foxp2 dosage on striatal circuits, a potential effect of the “humanized” knock-in allele on Foxp2 dosage in specific striatal regions and compartments is unknown. As RNA and protein expression levels are unchanged in *Foxp*^*hum/hum*^ when measured in larger tissue punches (Enard et al., 2009; Schreiweis et al., 2014), we argued that any dosage effect may be restricted to specific anatomical structures. To explore this hypothesis, we analyzed Foxp2 expression in three striatal subregions (ventral, dorsomedial and dorsolateral striatum) as well as striosome-matrix compartments of adult *Foxp2*^*hum/hum*^ mice and wildtype controls, and investigated whether there is a behavioural modification associated with potential histological changes (Figure 1B).

## MATERIALS AND METHODS

### Animals

*Foxp2*^*hum/hum*^ mice and wildtype littermates of the 5H11 line (C57Bl/6), as described in Enard et al., 2009, were housed in a 12 hour light-dark cycle with food and water available ad libitum. Experiments were performed in accordance with French (Ministère de l’Agriculture et de la Foret, 87-848) and European Economic Community (86-6091) guidelines for laboratory animal care and approved by the “Direction Départementale de la Protection des Populations de Paris” (license B75-05-22).

### Validation of specificity of the antibody against Foxp2

In order to test the specificity of the antibody against Foxp2 used in the histological quantification (goat polyclonal IgG, Santa Cruz sc-21069; validated previously also in (Vernes et al., 2011)), we performed a Western blot analysis using newborn mouse striata of wildtype mice and homozygous Foxp2 knock-out mice *(Foxp2*^*-/-*^), which are characterized by the absence of the Foxp2 protein (Groszer et al 2008) (Supplementary Figure S2). Briefly, *Foxp2*^*wt/wt*^ and *Foxp2*^*-/-*^ mouse pups, aged P0-P1, were decapitated, the striatum dissected in ice-cold PBS, snap-frozen in liquid nitrogen and stored at −80°C. Striatal tissue was homogenized either in 95°C hot solution of 1% SDS and 1mM sodium orthovanadate (Supplementary Figure S2, lanes 1-2), or in RIPA buffer with proteinase inhibitors (Supplementary Figure S2, lanes 3-4), pulled through a syringe and sonicated afterwards. Protein content of the lysates was determined using a Bicinchoninic-Assay (BCA). 40 μg of proteins were separated by SDS-PAGE (4-12% Bis-Tris NuPAGE gels, Invitrogen) for 30 min at 80 V and subsequently at 130 V for 1.5 hours before electrophoretic transfer onto PVDF membranes (Immobilon-P Membrane, PVDF, 0.45 μM). Gels were stained with Coomassie Blue and membranes stained with Ponceau Red to check for successful transfer. Membranes were blocked for 2 hours at room temperature in blocking buffer (1% non-fat milk TBST (0.075% Tween 20)) before being incubated overnight in blocking buffer with the anti-Foxp2 antibody (Foxp2 (N-16), sc-21069, Santa Cruz, dilution: 1:500) and anti-Actin antibody (Millipore, MAB1501, dilution: 1:5000). The incubated membranes were washed three times for 10 min in TBST at RT and incubated for 2 hours at room temperatured with secondary antibodies (Rockland, IRDye Goat anti-mouse 800nm; Donkey anti-Goat 800nm; dilution: 1:4000). Membranes were again washed 3x 10 min in TBS-T and imaged (Odyssey Imaging Systems).

### Histology

Animals homozygous for the human version of *Foxp2* and their wildtype siblings, aged 2.6 to 5.1 months, were anesthetized with an intraperitoneal injection of pentobarbital (200mg/kg), transcardially perfused with 30ml of 4°C cold 0.9% sodium chloride solution, followed by 60ml of 4°C cold 4% PFA in 1xPBS. Fixed brains were carefully dissected and postfixed in 4% PFA in 1x PBS overnight on a spinning wheel at 4°C, rapidly washed three times in 1xPBS and stored in 1x PBS with 0.025% sodium azide at 4°C until further processing. After dehydration in 15% sucrose in 1xPBS for 24 hours and in 30% sucrose for 48 hours, brains were embedded in O.C.T. compound and sectioned in ten series of 40μm on a cryostat (CryoStar NX70, Thermo Scientific). Sections were stored in 1xPBS with 0.025% sodium azide until immunofluorescent staining. Heat-induced epitope retrieval was performed for 20 minutes at 85°C in 2ml of 10mM sodium citrate buffer with 0.05% Tween 20 at pH 6.0 and allowed to cool down to room temperature for 30 minutes. After three washes for 5 minutes in ddH_2_0 followed by three washes of 10 minutes in 1xTris-buffered saline with 0.1% Tween 20, sections were blocked for three hours in 1x Tris-buffered saline with 0.1% Tween 20, 5% normal donkey serum and 1% BSA, on an orbital shaker at room temperature. Sections were incubated in the same blocking buffer with primary antibodies (anti-Foxp2: goat polyclonal IgG, Santa Cruz sc-21069, 1:1000; anti-MOR1: rabbit polyclonal IgG, Immunostar #24216, 1:500) at 4°C overnight on an orbital shaker. Subsequently, sections were washed three times for 10 minutes in 1x Tris-buffered saline with 0.1% Tween 20, incubated in blocking buffer with secondary antibodies at 4°C on an orbital shaker for three hours (dilution 1:500; Alexa Fluor Donkey anti-Goat 633 (Invitrogen A21082), Donkey anti-rabbit Cy3 (Jackson Immuno #711-165-152)), washed once for 40 minutes and twice for 10 minutes in 1x Tris-buffered saline with 0.1% Tween 20. A final 10 minute rinse in 1x Tris-buffered saline was performed and the sections mounted and coverslipped in fluorescence-protecting mounting medium (Vectashield H-1000). Mutant and wildtype mice were matched in comparable (age, sex) pairs, perfused and sectioned in a pseudorandom order for genotype. All sections were stained as one batch and the well position on the staining plates were also pseudorandomly balanced, correcting for any potential bias due to processing order or staining procedure. The experimenter was blind for the genotype during the entire sample treatment.

### Confocal Imaging and Histology Quantification

In 14 mice (n = 4 wildtype females and homozygous knock-in males; n = 3 wildtype males and homozygous knock-in females), the striatum at bregma level 0.86 was analyzed. Acquisition of the entire hemisphere was done using a LEICA SP5 confocal microscopy with a 40x objective. Five confocal planes at 1μm distance were analyzed using IMARIS 8.1 software (Bitplane), assuring unbiased, automatized quantification with constant acquisition parameters for all sections. We counted all nuclei that were Foxp2 immunofluorescence-positive, defined as spots of 9μm diameter. In order to illustrate the shift to higher expression values in ‘humanized’ mice, we chose a mean intensity threshold of 62 [intensity range 0-255] that we defined in four steps. Visually, Foxp2 expressing cells with an intensity level of or above 62 were perceived as a salient and separate population. This visual impression was confirmed via mean intensity distribution plots (Supplementary Figure S1B) as well as via cumulative distribution function analysis (Figure 2E, F). This showed that intensity level 62 is close to the middle of the intensity shift (Figure 2E, F; Supplementary Figure S1B) in the ventral striatum, the most prominent shifted region as verified via QQ-Plots (data not shown) and cumulative distribution function analysis (Figure 2F). The highly Foxp2 expressing cells were designated Foxp2^+ high^ and their number per volume was measured in three striatal subregions and the matrix as well as the striosome compartment. We distinguished between ventral, dorsomedial and dorsolateral striatal subregions. The dorsal regions were divided by a straight vertical line running through the midline of the dorsal striatum; the ventral striatum was defined as the striatum below a straight separation line running medially from the upper tip of the nucleus accumbens to the upper tip of the lateral stripe of the striatum laterally. All striosomes and three matrix compartments were defined as MOR1+ and MOR1-compartments, respectively, and manually outlined as regions of interest. Images were acquired in a pseudorandom fashion across matched pairs to exclude any genotype bias during the imaging and quantification procedure; the experimenter was blinded for the genotype during the entire imaging and quantification process.

**Figure 2.**
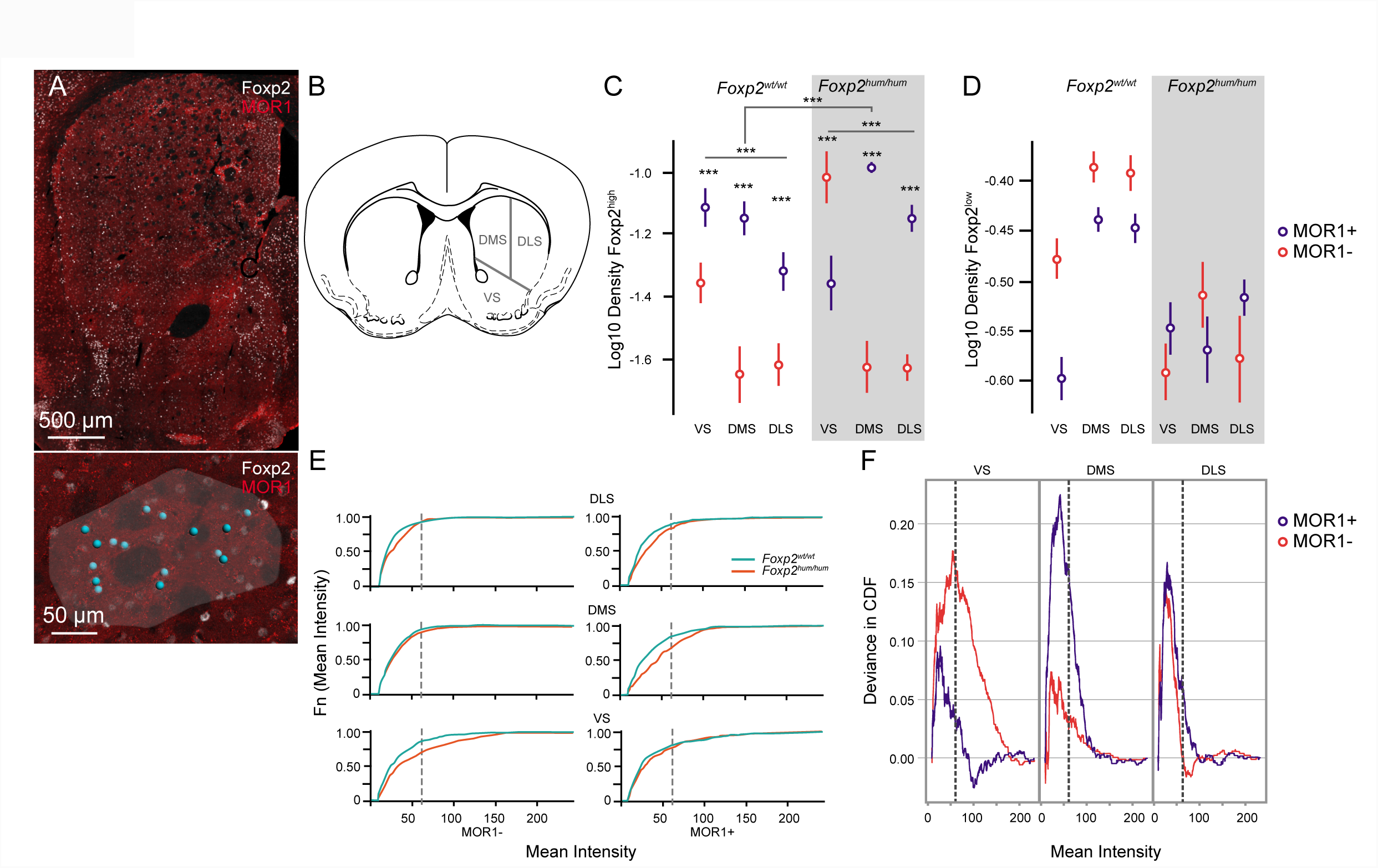
Humanized Foxp2 shifts striatal Foxp2 expression in a region- and compartment-specific manner. (A) Coronal section of the adult mouse striatum, immunohistochemically labelled for Foxp2 (white) and μ opioid receptor 1 (MOR1, red) (upper panel). Magnification of the same section with an exemplary, MOR1 + striosome (highlighted in a transparent white); automated counts of highly Foxp2 expressing signals are illustrated in blue (lower panel). (B) Illustration of representative coronal mouse brain section, modified from (Paxinos & Franklin, 2001), illustrating the subregional division into ventral (VS), dorsomedial (DMS), and dorsolateral striatum (DLS). (C) Average logarithmic density of highly Foxp2 expressing cells, corrected for pseudoreplication, in VS, DMS and DLS in MOR1- (red) and MOR1+ (blue) compartments. Error bars represent +/- SEM. ***p<0.001. (D) Average logarithmic density of weakly Foxp2 expressing cells, corrected for pseudoreplication, in VS, DMS and DLS in MOR1- (red) and MOR1+ (blue) compartments. Logarithms of weakly Foxp2 expressing densities could not be statistically modelled. (E) Empirical cumulative distribution function of mean Foxp2 intensity values, stratified by region (VS, DMS, DLS), compartment (MOR1-on the left, MOR1+ on the right of each graph) and genotype (*Foxp2*^*hum/hum*^ in orange; *Foxp2*^*wt/wt*^ in turquoise). (F) Deviance in cumulative distribution functions (CDF) between humanized Foxp2 and wildtype mice, according to striatal subregion (VS, DMS, DLS) and compartment (MOR1-: red; MOR1+: blue).

### Locomotor sensitization assay

Locomotor activity was measured in a circular corridor, equipped with four infrared beams placed every 90° (Imetronic, France). Counts were incremented by consecutive interruption of two adjacent photobeams, i.e. mice passing a quarter of the corridor. After an initial saline injection, mice were placed in the activity corridors for 60 minutes, followed by a subcutaneaous injection of either 0.9% NaCl solution at 0.1ml/10g body weight (on each of the three consecutive habituation days), or 10 mg/kg morphine diluted in 0.9% NaCl solution at the same volume/weight ratio (on the five consecutive days of acute morphine treatment and morphine sensitization), and their locomotor activity was monitored during 180 minutes post injection. The experimenter was blind for the genotype during experimentation. Injections of mutant and wildtype pairs were ordered in a pseudorandom manner across all pairs and pseudorandomly assigned to a position in the apparatus, which stayed constant during the entire treatment.

### Statistical analysis

Data analysis was performed using R and its packages “tidyverse”, “lme4”, “lmerTest”. Foxp2 densities were log10-transformed, visually inspected for normal distribution using histogram, Q-Q and density plot and a Shapiro-Wilk Normality Test was performed. All statistical models were corrected for repeated measurements per animal.

For Foxp2+ densities, a linear mixed effect model with log10 densities as response variable was fitted with fixed effects *sex, genotype, brain region* and *compartment* and random effects with an intercept for animal identity to account for individual mouse differences and repeated measurements of Foxp2+ signal within the same mouse. Model selection was based on comparison of AIC (Akaike Information Criterion), BIC (Bayesian Information Criterion) and LRT (Likelihood Ratio Test) of nested models. Model assumptions (normality, homoscedasticity, outliers) were visually inspected using model residuals. The explanatory variables were tested for significance based on restricted maximum likelihood estimation (REML) and the Satterthwaite’s method for approximating degrees of freedom for the F tests.

For the analysis of morphine sensitization, a generalized linear mixed effect model with movement rate as response variable assuming a negative binomial distribution was fitted with fixed effects including treatment and genotypes as well as polynomial coding for day and time point to account for dependency of temporal patterns. The individual time points were replaced by time windows: baseline given by the mean movement rate over 60 minutes prior to recording, 1 to 30 minutes as initial phase, 40 to 90 as plateau phase and 100 to 140 as end phase.

### Foxp2 nomenclature

Following standard nomenclature for FOXP2 is used in this text: *italics* font refers to the gene and roman font to the protein. Upper case letters refers to the human version (FOXP2); upper case F but otherwise lower case letters refer to the murine version (Foxp2); and upper case F and P but otherwise lower case letters refer to any other variants of FoxP2, e.g. the avian or chimpanzee version.

## RESULTS

To assess patterns of Foxp2 protein expression in adult striatal tissue, we used an unbiased, semi-automated approach. We first quantified all Foxp2+ nuclei across all detectable levels of staining intensities (Fig.2, Supplementary Fig. 1). We found that *Foxp2*^*um/hum*^ mice show a shift to higher staining intensities (Fig.2E; Supplementary Figure S1B). This shift was not uniform throughout the striatum but rather pronounced in specific regions and compartments (Figure 2E, F; Supplementary Figure S1B-G; interaction effect *genotype, region, compartment: F*_*2*_, _7548_ = 21.34; *p*=0; main effect *genotype: F*_*1,11*_ *=* 1.36, *p=*0.27; Table 1). To statistically analyse and simply illustrate this shift, we chose an intensity cut-off (see methods). All Foxp2+ nuclei above this cut-off were designated Foxp2^high^ and their number per tissue volume (=density) quantified (n=7 mice/group). In each of three striatal subregions (ventral (VS), dorsomedial (DMS) and dorsolateral striatum (DLS), we distinguished the striosome and the matrix by MOR+ and MOR-staining respectively and quantified Foxp2^high^ nuclei in 3-6 volumes per compartment, region and mouse (Figure 1B; Figure 2A-B). We used the logarithm of the Foxp2^high^ density as response variable to analyse the effects of the predictive variables *sex, region, compartment* and *genotype* by a linear mixed effect model that allows taking into account the correlation of several measurements per mouse. We find that the use of a linear mixed effects model is warranted as individual variation of mice/section explains 33% of the total variation and as the model assumptions (i.e. normal and homoscedastic residuals without strong outliers) are not violated (not shown). Using standard approaches for model selection, we continued to analyse the effects of *region, compartment and genotype*, controlling for *sex*. We find that while *sex* has no significant effect on Foxp2^high^ densities, *region, compartment* and *genotype* do (Table 2). Concordant with previous data that showed enrichment of *Foxp2* RNA in developing striosomes of primates (Takahashi et al., 2003, 2008) and of Foxp2 protein in developing striosomes of mice (Chen et al., 2016), we also find a higher number of Foxp2^high^ nuclei in adult striosomes in all three regions of wildtype mice (Figure 1C; main effect compartment: *F*_*1,202*_= 51.53; *p=*0). Furthermore Foxp2^high^ density is higher in the ventral striatum than in the two dorsal regions (Fig 2C; main effect *region: F*_*2,202*_ *=* 9.84; p<0.0001). Importantly, the presence of the humanized version of *Foxp2* significantly influences this distribution in a compartment and region-specific manner (Fig. 2C-F; Supplementary Figure S1B, E, F; interaction effect *genotype, region, compartment F*_*2,202*_ *=* 8.19; p=0.0004). While in the dorsal striatum the Foxp2^high^ density is increased ∼ 2-fold in *Foxp2*^*hum/hum*^ mice only in striosomes, in the ventral striatum this was found for the matrix. Hence, the humanized Foxp2 does increase the density of highly Foxp2 positive neurons in the striatum in a region and compartment-specific manner, although detailed mechanisms of this effect will need to be explored in future studies applying methods such as ChIP Seq and single cell RNA Seq.

**Table 1:** Results of Generalized Linear Mixed Model analysis performed on log mean Foxp2 intensity values in adult striatal tissue

| *Predictor terms<sup>a</sup>* | Sum Sq | Mean Sq | NumDF | DenDF | F value | Pr(>F) |
| --- | --- | --- | --- | --- | --- | --- |
| *genotype<sup>a</sup>* | 0.5623346 | 0.5623346 | 1 | 11.00233 | 1.3588318 | 0.2683889 |
| *region<sup>a</sup>* | 35.4613682 | 17.7306841 | 2 | 7547.84426 | 42.8446270 | 0.0000000 |
| *compartment<sup>a</sup>* | 39.6027268 | 39.6027268 | 1 | 7547.62627 | 95.6964801 | 0.0000000 |
| *genotype :region<sup>a</sup>* | 0.1140572 | 0.0570286 | 2 | 7547.84426 | 0.1378046 | 0.8712711 |
| *genotype:compartment<sup>a</sup>* | 0.2371118 | 0.2371118 | 1 | 7547.62627 | 0.5729597 | 0.4491088 |
| *region:compartment<sup>a</sup>* | 27.9239318 | 13.9619659 | 2 | 7547.84426 | 33.7378535 | 0.0000000 |
| *genotype :region :compartment<sup>a</sup>* | 17.6602460 | 8.8301230 | 2 | 7547.84426 | 21.3372098 | 0.0000000 |

<sup>a</sup>Predictor variables and their interaction terms of the final statistical model. Interactions are marked by ':' between the names of the predictors. The levels of each predictor are as follows: sex (male, female), region (ventral, dorsomedial, dorsolateral striatum), compartment (striosome, matrix), genotype (hum/hum, wt/wt).

**Table 2:** Results of Generalized Linear Mixed Model analysis performed on the Foxp2+ high quantification in adult striatal tissue

| *Predictor terms<sup>a</sup>* | Sum Sq | Mean Sq | NumDF | DenDF | F value | Pr(>F) |
| --- | --- | --- | --- | --- | --- | --- |
| *sex<sup>a</sup>* | 0.0070031 | 0.0070031 | 1 | 10.00643 | 0.0711388 | 0.7951017 |
| *region<sup>a</sup>* | 1.9377606 | 0.9688803 | 2 | 202.15019 | 9.8420339 | 0.0000834 |
| *compartment<sup>a</sup>* | 5.0723474 | 5.0723474 | 1 | 202.07428 | 51.5256779 | 0.0000000 |
| *genotype<sup>a</sup>* | 0.0301606 | 0.0301606 | 1 | 10.01806 | 0.3063760 | 0.5920505 |
| *region:compartment<sup>a</sup>* | 3.8504327 | 1.9252163 | 2 | 202.06706 | 19.5566409 | 0.0000000 |
| *region:genotype<sup>a</sup>* | 0.0066591 | 0.0033295 | 2 | 202.16369 | 0.0338219 | 0.9667491 |
| *compartment:genotype<sup>a</sup>* | 0.0893297 | 0.0893297 | 1 | 202.07965 | 0.9074244 | 0.3419367 |
| *region:compartment:genotype<sup>a</sup>* | 1.6126950 | 0.8063475 | 2 | 202.06396 | 8.1910007 | 0.0003798 |

<sup>a</sup>Predictor variables and their interaction terms of the final statistical model. Interactions are marked by ‘:’ between the names of the predictors. The levels of each predictor are as follows: sex (male, female), region (ventral, dorsomedial, dorsolateral striatum), compartment (striosome, matrix), genotype (hum/hum, wt/wt).

The increased density of Foxp2^high^ nuclei was surprising, as an overall increase in Foxp2 protein or RNA levels in *Foxp2*^*hum/hum*^ mice had not been observed previously (Enard et al., 2009; Schreiweis et al., 2014). As our threshold for Foxp2 staining was rather conservative, we next also analysed the density of Foxp2 stained nuclei that fell below the defined thresholds. Unfortunately, analysing the data using linear effects models was not possible as negative deviance values indicated unstable model fitting. However, the plotting of mean values (Fig. 2D; Supplementary Figure S1F) and the plotting of intensity distributions (Fig. 2E; Supplementary Figure S1B) clearly indicated that neurons weakly stained for Foxp2 (Foxp2^low^) were also affected by the humanized *Foxp2* allele, but in a reverse manner than strongly stained Foxp2 (Foxp2^high^) neurons. In general, there was a decrease in Foxp2^low^ expression in all regions and compartments. Hence, the total amount of Foxp2 protein does not seem to be affected by *Foxp2*^*hum*^ (main effect *genotype: F*_*1,11*_*-*1.36, *p=*0.27; Supplementary Figure S1C), in agreement with previous observation on total striatal Foxp2 concentrations (Enard et al., 2009). Instead, it seems that *Foxp2*^*hum*^ leads to a shift towards higher Foxp2 expression in medium spiny neurons in a region and compartment-specific manner. The observed changes in Foxp2 expression were not accompanied by changes in overall MOR1+ density, which corresponded to the densities reported in the literature (Johnston, Gerfen, Haber, & van der Kooy, 1990). This was true for the entire striatum (Wilcoxon-Mann-Whitney: *W* = 21, *p* = 1), as well as striatal subregions (Wilcoxon-Mann-Whitney: ***W*_*vs*_** =30, *p =* 0.23; ***W*_*DS*_** *=* 12, *p =* 0.23) (Supplementary Figure 3).

To test whether the observed histological change of humanized *Foxp2* is associated with a modification of behaviour, we chose a morphine sensitization assay. Morphine stimulates the µ opioid receptors (MOR) that are strongly enriched in striosomes, and is thought to involve a cross-reaction of dopaminergic and opioid brain systems (Badiani, Belin, Epstein, Calu, & Shaham, 2011). We injected 13 female *Foxp2*^*hum/hum*^ mice and 10 female wildtype controls with saline for three consecutive days and then with morphine for five consecutive days. We found that morphine induced strong locomotion compared to saline injections (Figure 3) as described before (Babbini & Davis, 1972). There is a tendency for *Foxp2*^*hum/hum*^ mice to move less after morphine injection that is significant on day 2 (Wilcoxon Rank Test, p= 0.02). In addition, we performed a linear mixed effect model to take into account the correlation of several measurements per mouse. To that end, we used a quartic polynomial regression to model the movement after morphine injection in the initial phase (0-30 minutes after morphine injection), plateau phase (40-90 min) and end phase (100-140min) across days. We did not detect any difference in locomotion between *Foxp2*^*hum/hum*^ and wildtype mice during the acute morphine treatment on day 1 (Wald T-test, *p*_*Genotype * Treatment*_ *=0*.24; Table 3; Figure 2, central panel). However, we observed a tendency for an interaction of *timepoint* (linear component) and *genotype* during the sensitized morphine treatment of day 2-5 (Wald T-test; *P*_*TimePoint*.*LGenotype*_*=*0.04; Table 4; Figure 2, right panel). Taken together, our results suggest that humanized Foxp2 impairs striatal-dependent, morphine-evoked behavioural adaptations.

**Figure 3.**
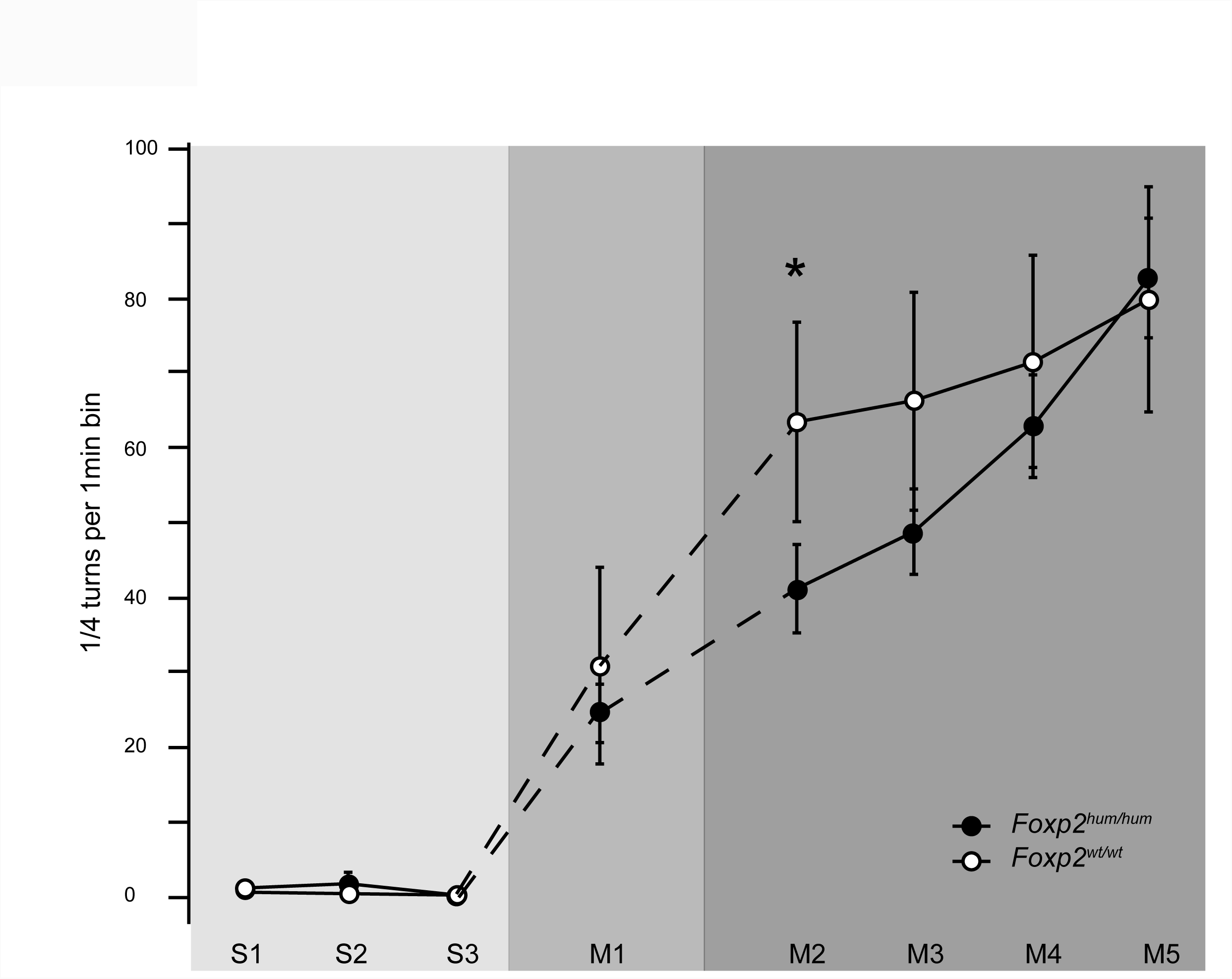
Influence of humanized Foxp2 on morphine-induced locomotion. Average plateau of horizontal locomotion between 40 to 90 minutes after injection with 0.9% NaCl for three days (S1-3, left panel), after acute (M1) or chronic (M2-5) injection with morphine, diluted in 0.9% NaCI (lOmg/kg, s.c.) of humanized (filled dots) compared to wildtype mice (white dots), +/- SEM. Each data point represents one of eight subsequent days of treatment. *: p<0.05.

**Table 3:** Results of Generalized Linear Mixed Model analysis on acute morphine treatment (10mg/kg)

| *Predictor terms<sup>a</sup>* | estimate | std.error | statistic | p.value | group |
| --- | --- | --- | --- | --- | --- |
| *Intercept<sup>a</sup>* | 0.9423235 | 0.2241036 | 4.2048557 | 0.0000261 | fixed |
| *TimePoint.L<sup>a</sup>* | -0.6854785 | 0.2142677 | -3.1991682 | 0.0013782 | fixed |
| *TimePoint.Q<sup>a</sup>* | 0.1607373 | 0.1862741 | 0.8629071 | 0.3881885 | fixed |
| *TimePoint.C<sup>a</sup>* | -0.2277128 | 0.1753738 | -1.2984425 | 0.1941353 | fixed |
| *Treatment<sup>a</sup>* | 1.9296549 | 0.3101640 | 6.2214023 | 0.0000000 | fixed |
| *Genotype<sup>a</sup>* | -0.3908863 | 0.2879686 | -1.3573919 | 0.1746567 | fixed |
| *Treatment:Genotype<sup>a</sup>* | 0.4386409 | 0.3724600 | 1.1776858 | 0.2389219 | fixed |

<sup>a</sup>Predictor variables and their interaction terms of the final statistical model. Interactions are marked by ‘:’ between the names of the predictors.

**Table 4:** Results of Generalized Linear Mixed Model analysis on chronic morphine treatment (days 2-5; 10mg/kg)

| *Predictor terms<sup>a</sup>* | estimate | std.error | statistic | p.value | group |
| --- | --- | --- | --- | --- | --- |
| *Intercept<sup>a</sup>* | 2.9648968 | 0.1163563 | 25.4811873 | 0.0000000 | fixed |
| *Day.L<sup>a</sup>* | 0.1124145 | 0.0507264 | 2.2160958 | 0.0266849 | fixed |
| *Day.Q<sup>a</sup>* | 0.1366523 | 0.0505371 | 2.7040003 | 0.0068510 | fixed |
| *Day.C<sup>a</sup>* | -0.0160882 | 0.0506218 | -0.3178122 | 0.7506274 | fixed |
| *TimePoint.L<sup>a</sup>* | 1.5521370 | 0.0910098 | 17.0546074 | 0.0000000 | fixed |
| *TimePoint.Q<sup>a</sup>* | -1.5972004 | 0.0819931 | -19.4796918 | 0.0000000 | fixed |
| *TimePoint.C<sup>a</sup>* | -0.0322521 | 0.0723235 | -0.4459422 | 0.6556390 | fixed |
| *Genotype<sup>a</sup>* | -0.0365738 | 0.1549870 | -0.2359795 | 0.8134486 | fixed |
| *TimePoint.L:Genotype<sup>a</sup>* | 0.2532755 | 0.1233332 | 2.0535870 | 0.0400157 | fixed |
| *TimePoint.Q:Genotype<sup>a</sup>* | 0.0917688 | 0.1106434 | 0.8294105 | 0.4068721 | fixed |
| *TimePoint.C:Genotype<sup>a</sup>* | 0.1084997 | 0.0964801 | 1.1245810 | 0.2607666 | fixed |

<sup>a</sup>Predictor variables and their interaction terms of the final statistical model. Interactions are marked by ‘:’ between the names of the predictors.

## DISCUSSION

We have found that striatal Foxp2 expression levels in medium spiny neurons are shifted in mice homozygous for the humanized allele of *Foxp2*. Several findings indicate that such a shift could functionally matter. In general, Foxp2 expression levels are associated with functional consequences. Foremost, haploinsufficiency is thought to be the major etiological factor for phenotypes observed in humans with genetic disruption of one *FOXP2* allele (Vargha-Khadem et al., 2005). Additionally, D1R+ and D2R+ medium spiny neurons associate with higher and lower Foxp2 expression levels, respectively (Vernes et al., 2011). Foxp2 expression changes during development (Takahashi et al., 2003, 2008) and, dependent on Foxp1 expression levels, the identity of motor neurons in the brainstem is specified (Dasen, De Camilli, Wang, Tucker, & Jessell, 2008; McDole et al., 2018; Rousso, Gaber, Wellik, Morrisey, & Novitch, 2008). In songbirds, FoxP2 expression levels in medium spiny neurons of Area X are associated with different learning stages (Thompson et al., 2013), and FoxP2 knockdown interferes with song learning (Haesler et al., 2007; Murugan et al., 2013) mediated by D1R-dependent modulation of corticostriatal activity (Murugan et al., 2013). Given these findings, it is plausible that our observed shift in Foxp2 expression levels in *Foxp2*^*hum/hum*^ mice could be directly functionally relevant.

That this shift is highly region- and compartment-specific explains why no differences in average Foxp2 expression levels have been seen in *Foxp2*^*um/hum*^ mice previously. Furthermore, as striatal subregions are differently recruited during different stages of learning (Graybiel, 2008; Miyachi, Hikosaka, & Lu, 2002; Thorn, Atallah, Howe, & Graybiel, 2010; Yin et al., 2009) and the striosome-matrix compartments also differ in terms of development, connectivity, and function (Bloem et al., 2017; Brimblecombe & Cragg, 2015; Crittenden & Graybiel, 2011; Gerfen & Bolam, 2010; Yoshizawa et al., 2018), such a region- and compartment-specific shift could have specific consequences. This is in line with our previous region-specific effects in *Foxp2*^*um/hum*^ mice with respect to dopamine levels, synaptic plasticity and learning behaviour (Schreiweis et al., 2014) and could also be related to the behavioural changes in the morphine sensitization assay studied here. The mechanisms and cause-effect relationships among those phenotypes in *Foxp2*^*um/hum*^ mice must be further studied in the future. In particular, it will be important to disentangle whether these phenotypes are set up during development or can also be caused when Foxp2 is humanized only in adults. Any effect of the humanized allele on developmental wiring of the striatum may affect expression patterns, as Foxp2 levels differ in striatal neurons that project to different target regions (Gokce et al., 2016; Vernes et al., 2011), or vary with neuronal activity (Horng et al., 2009; Teramitsu & White, 2006). Furthermore, it will be crucial to determine in which cells humanized Foxp2 must be present to cause these effects. This will be important to interpret molecular data generated using e.g. single-cell RNA-sequencing.

To what extent the striatal region-specific findings in *Foxp2*^*um/hum*^ mice reflect changes that occurred during human evolution is naturally more speculative and much less amenable to direct experimentation. However, in this context it is interesting to note that humans lacking one copy of functional FOXP2 show different neuroanatomical and neurofunctional effects in the caudate nucleus and the putamen - thought to be the primate homologues of the rodent DMS and DLS, respectively (Liégeois et al., 2003; Vargha-Khadem et al., 1998). Thus, our findings could reflect a contribution of two amino acid changes in FOXP2 towards circuit-specific evolutionary changes in the striatum that have been recruited and adapted for speech and language development. While certainly not the only contribution to the evolution of speech and language, studying the human-specific properties of FOXP2 opens new avenues for dissecting mechanisms underlying this human trait.

## CONTRIBUTIONS

LL and CS developed the histology protocol and the Western Blotting protocol. LL and CS performed Western Blotting. CS and FO perfused mice for the histology study. CS conceptualized and conducted all histology and sensitization experiments. TI imaged the histology samples and performed the image analysis and quantifications. CS, TI, BV, MG and WE analysed the data. CS, TI, BV, WE and MG wrote the manuscript. CS conceptualized, and TI, BV and CS produced the figures. MG supervised experiments. CS, TI, BV, LL, FO, EB, WE and MG edited the manuscript.

## ACKNOWLEDGEMENTS

The authors thank Ines Bliesener and Mythili Savariradjane for excellent technical assistance, Dr. Thomas Lemmonier for advice on immunofluorescence staining, Dr. Karim N’Diaye for his important help during the initial statistical analyses, and Prof Svante Pääbo for generously providing *Foxp2*^*hum/hum*^ mice and scientific mentorship.

---

**Supplementary Figure 1.**
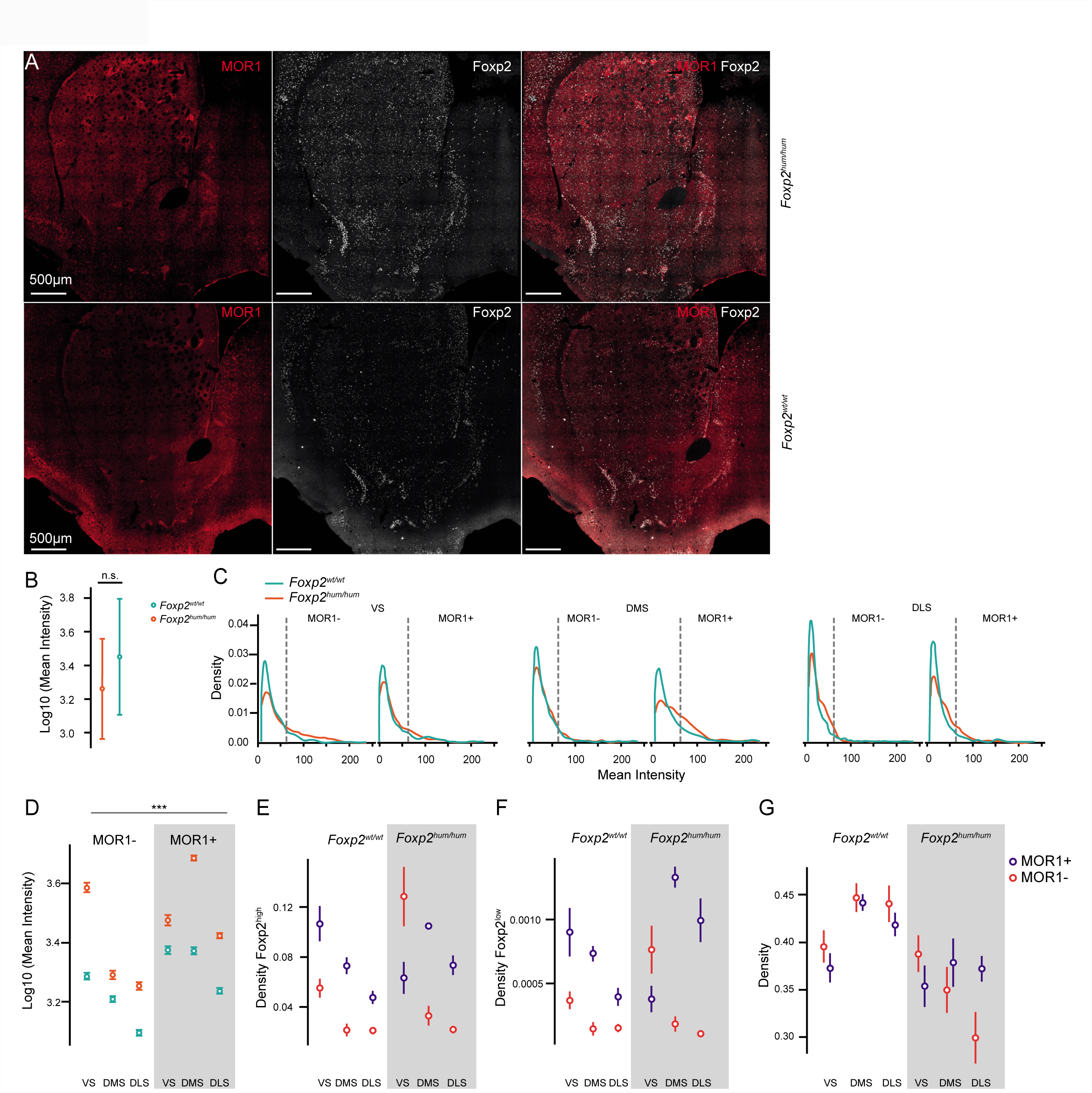
Immunofluorescent staining and quantification. (A) Examples of MOR1 (red), Foxp2 (white) and MOR1/Foxp2 overlay images of analysed coronal sections to illustrate the observed difference in Foxp2 intensity distribution. Upper panels represent a *Foxp2*^*hum/hum*^, lower panels represent a wildtype mouse. (B) Log-10 of mean Foxp2 intensities showed no overall genotype effect *(genotype*: *F*_*1,11*_=1.36, *p*=0.27; *Foxp2*^*hum/hum*^ in orange; *Foxp2*^*wt/wt*^ in turquoise). Error bars indicate +/- SEM. (C) Density of mean Foxp2 intensity values, stratified by region (VS, DMS, DLS), compartment (MOR1-on the left, MOR1+ on the right of each graph) and genotype *(Foxp2*^*hum/hum*^ in orange; *Foxp2*^*wt/wt*^ in turquoise). The chosen cut-off for highly Foxp2 expressing cells at intensity level 62 is illustrated as a dashed line. (D) Log-10 of mean intensities, stratified by region (VS, DMS, DLS) and compartment (MOR1+, MOR1-) show a significant interaction of genotype*region*compartment (interaction effect *genotype, region, compartment:*. *F*_*2*,_,_7548_= 21.34; *p=*0; Table 1). (E) Average raw density of highly Foxp2 expressing cells, corrected for pseudoreplication, in VS, DMS and DLS in MOR1- (red) and MOR1+ (blue) compartments. (F) Average raw density of weakly Foxp2 expressing cells, corrected for pseudoreplication, in VS, DMS and DLS in MOR1- (red) and MOR1+ (blue) compartments. (G) Average raw densities of all Foxp2 expressing cells, stratified by region (VS, DMS, DLS), compartment (MOR1+, MOR1-), and genotype (*Foxp2*^*wt/wt*^, *Foxp2*^*hum/hum*^). Raw Foxp2 expressing densities could not be statistically modelled.

**Supplementary Figure 2.**
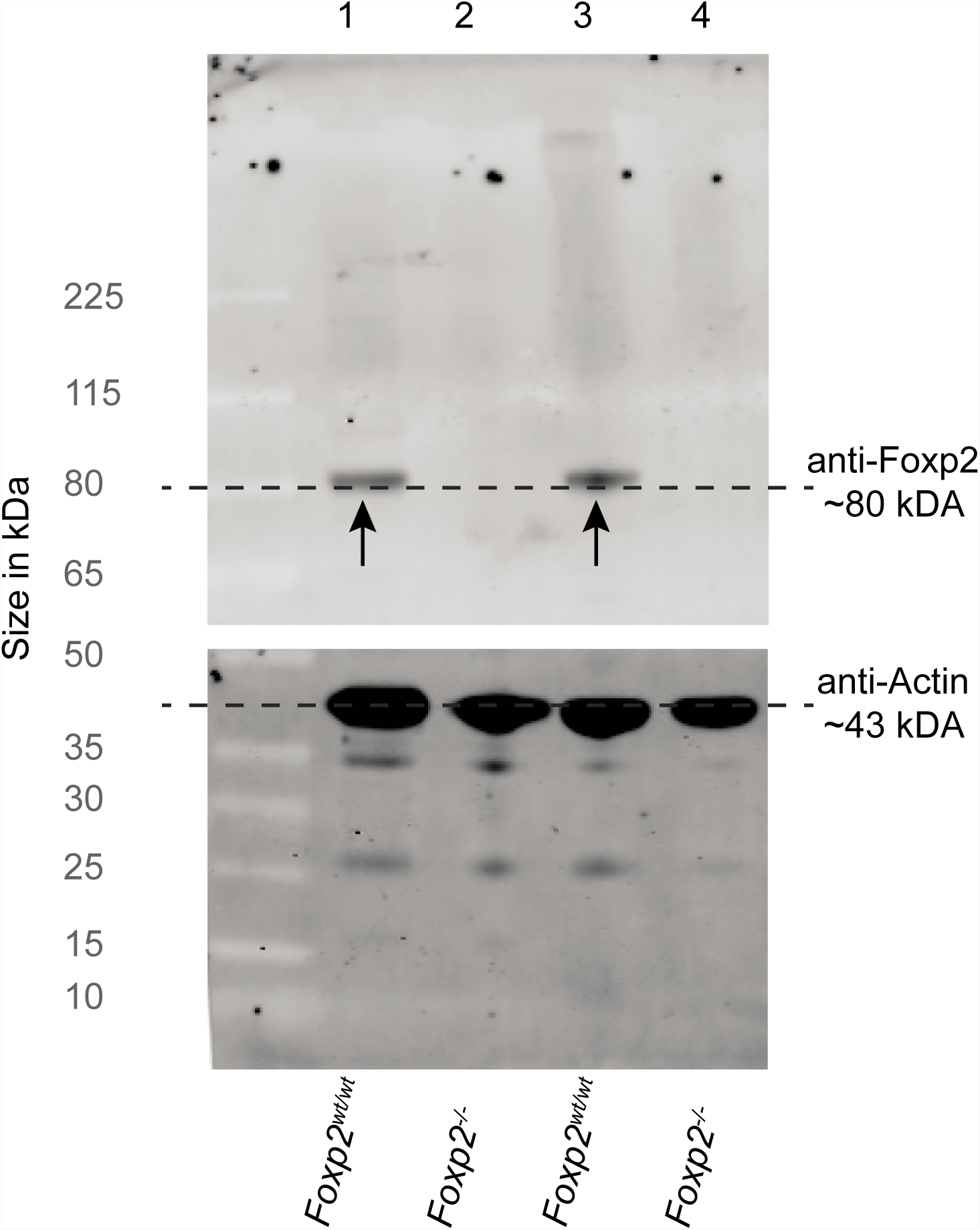
Validation of Foxp2 antibody specificity via Western Blot Analysis. Western Blot of whole-striatum protein of newborn wildtype (lanes 1, 3) and Foxp2 knockout mice, characterized by an absence of the Foxp2 protein (lanes 2, 4). Two different extraction protocols were used (see methods). The upper strip of the Western Blot was incubated with the Foxp2 antibody used in this study (dilution 1:500; goat polyclonal IgG, Santa Cruz sc-21069; previously also shown in (Vernes et al., 2011)); the lower strip was stained with anti-Actin antibody (Millipore, MAB1501, dilution: 1:5000) as a control. Grey dashed lines indicate the expected migration range for the 80kDA protein Foxp2 and approximately 43 kDA of the actin protein. Arrows indicate the band detected at 80kDA, corresponding to the expected band size of the Foxp2 protein.

**Supplementary Figure 3.**
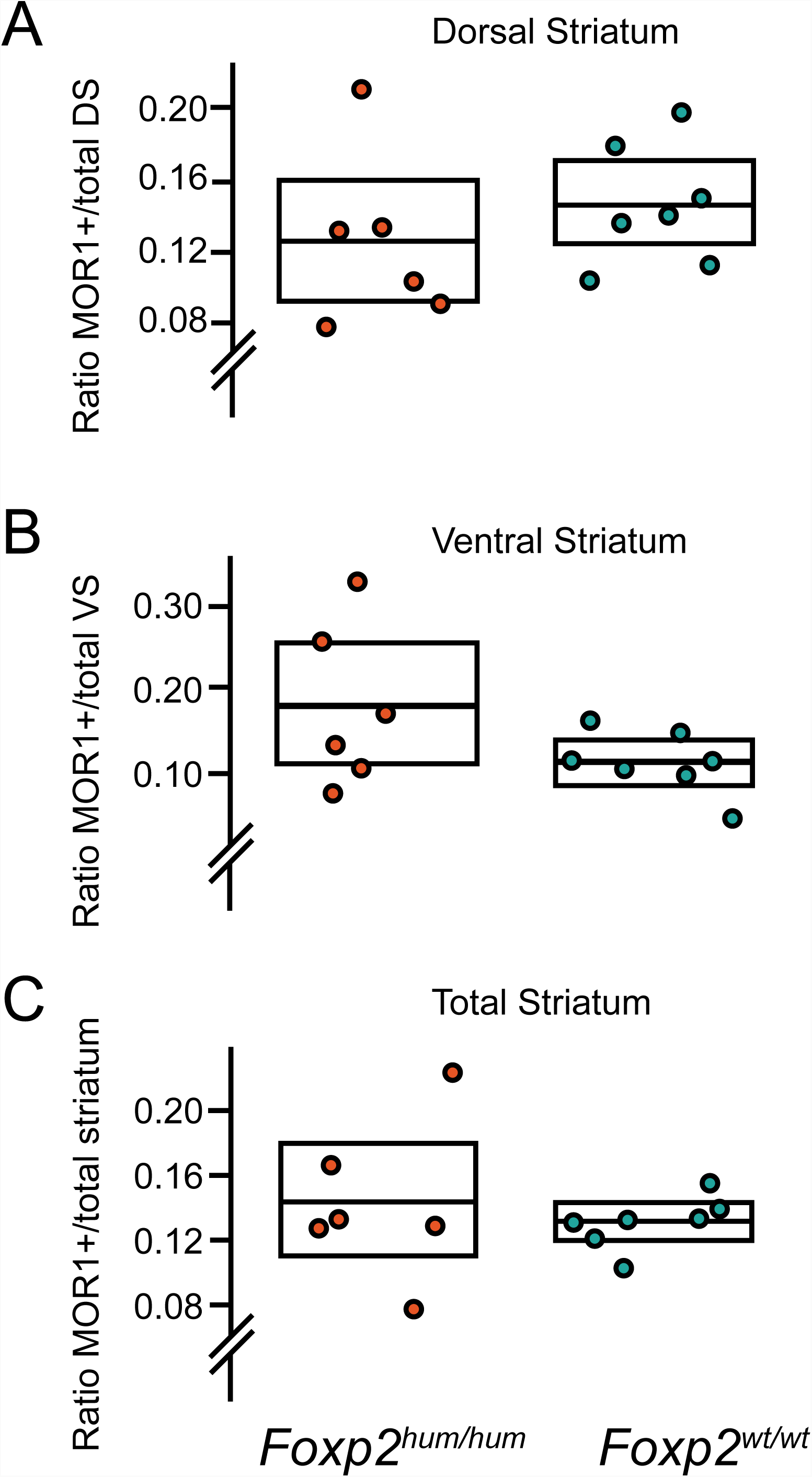
No major changes in striatal MOR1+ compartment structure in humanized Foxp2 mice. Box plots (indicated value is the median) showing the relative MOR1+ volume in the dorsal striatum (A), the ventral striatum (B), and the whole striatum (C), measured in the same sections in which we quantified Foxp2 expression. Individual points correspond to individual subjects. The measured 15% ratio of MOR1+ volume in the striatum corresponds to previously measured volumes (Johnston et al., 1990).

**Supplementary Figure 4.**
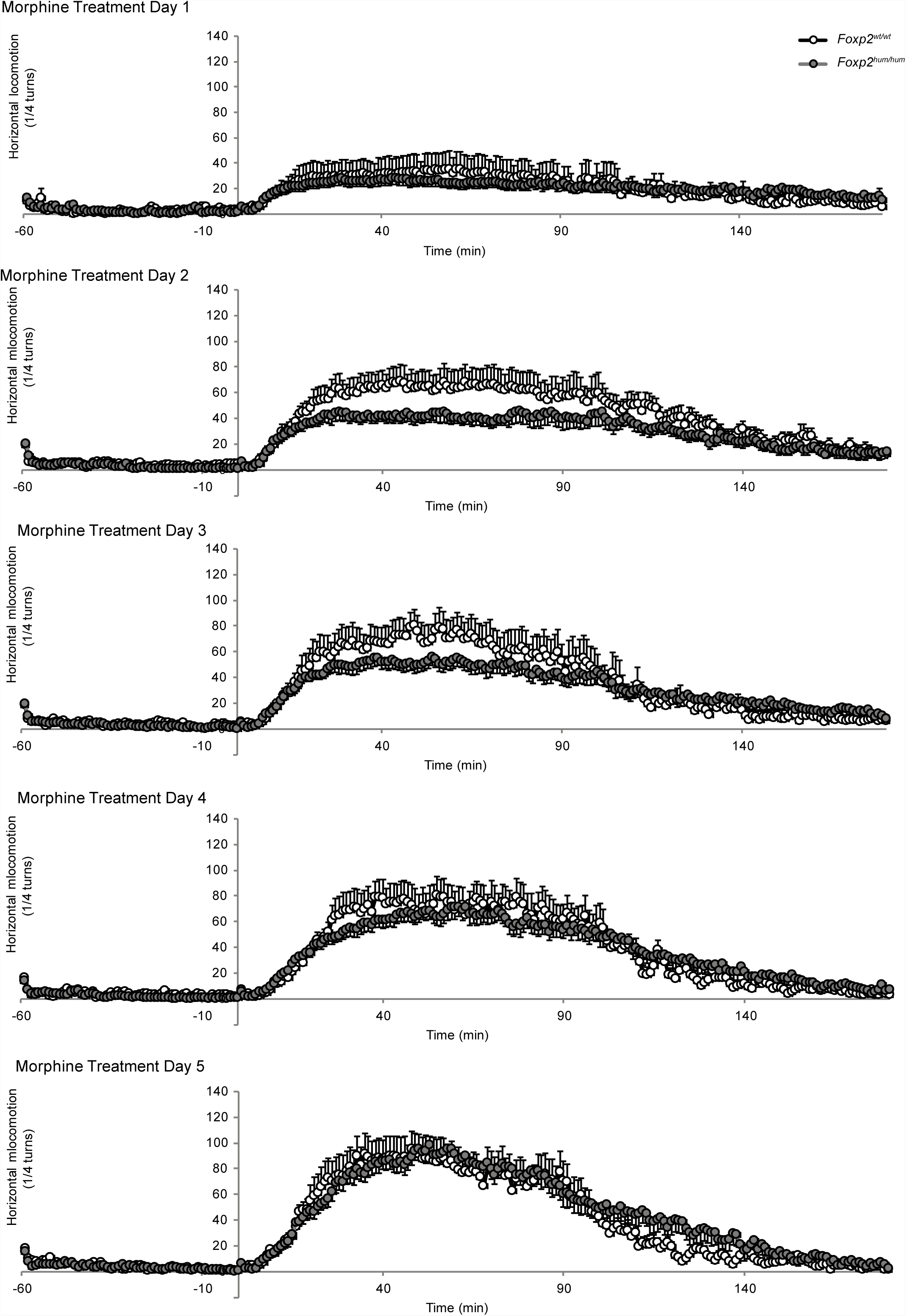
Comprehensive plots of horizontal locomotion under morphine treatment in humanized Foxp2 and wildtype mice. Individual plots of horizontal locomotion, measured as ¼ turns in the circular apparatus, in 1-minute bins from 60 minutes prior until 180 minutes after injection of morphine (lOmg/kg, s.c., diluted in 0.9% NaCl), in *Foxp2* ^*hum/hum*^ (grey dots) and *Foxp2*^*wt/wt*^ mice (white dots), +/- SEM.

